# Astroglial atrophy associates with loss of nuclear S100A10 in the hippocampus of aged male tree shrews

**DOI:** 10.1101/2024.07.04.602113

**Authors:** Juan D. Rodriguez-Callejas, Mariangel Irene-Fierro, Sara Aguilar-Navarro, Eberhard Fuchs, Claudia Perez-Cruz

## Abstract

Astrocytes are glial cells that participate in multiple physiological functions, such as protecting neurons against all types of damage. However, astrocytes develop morphological alterations during aging, such as decreased length, volume, and branch points of their processes, also known as astrocytic atrophy. Until now, the exact mechanism associated with the onset of atrophy is unknown. Tree shrew (*Tupaia belangeri*) is a long-lived animal from the Scadentia order that develops several age-dependent brain alterations. In this study, we analyzed the morphology of GFAP+ astrocytes in the hippocampal region of adult, old, and aged male tree shrews. Aged animals presented more GFAP+ astrocytes in the proximal subiculum, CA3, and CA2-CA1 subregions than younger animals, being significantly higher only in CA3. However, in aged subjects, the number of atrophic astrocytes was significantly higher in the dentate gyrus and CA2 subregion compared to old animals. Interestingly, in the proximal subiculum, astrocytes had a reduced arborization at all ages evaluated. S100A10, a protein overexpressed by neuroprotective-type astrocytes, was mainly found in the nucleus of astrocytes of adult and old subjects. However, in old and aged animals, S100A10 was located in the cytoplasmic compartment of atrophic astrocytes. Furthermore, in astrocytes of aged tree shrews, cytoplasmic inclusions of S100A10 colocalized with the nuclear export protein, Crm-1. These results suggest that the transport of S100A10 from the nucleus to the cytoplasmic compartment of astrocytes could be a process related to astroglial atrophy during the aging process in the hippocampus of three shrews.

## 1. Introduction

Astrocytes are present in all brain regions (Carson et al., 2006) and contribute to multiple physiological processes, such as regulation of the blood-brain barrier (BBB) (Clark & Mobbs, 1992; Janzer & Raff, 1987), secretion of neurotransmitters, cytokines, and neurotrophic factors (Araque et al., 1998; Condorelli et al., 1995; Liberto et al., 2004), uptake of neurotransmitters and ions (Anderson & Swanson, 2000; Karwoski et al., 1989; Liberto et al., 2004; Volterra & Steinhäuser, 2004), formation, modulation and elimination of synapses (W. Chung et al., 2015; W. S. Chung et al., 2013; Nedergaard, 1994; Parpura et al., 1994; Volterra & Steinhäuser, 2004), and provision of energy metabolites to neurons (Alberini et al., 2018; Brown & Ransom, 2007). Astrocytes also respond to all forms of brain damage via *reactive astrogliosis*. Reactive astrogliosis refers to diverse genetic, cellular, and functional alterations occurring in astrocytes, and it varies according to the intensity of the damage (Sofroniew & Vinters, 2010). Astrocytic hypertrophy is characterized by an enlargement of the cytoplasm and its processes, accompanied by overexpression of proteins that conform the intermediate filament system, such as the glial fibrillar acidic protein (GFAP). GFAP is a class III intermediate filament protein encoded by a gene in chromosome 17q21 (Garwood et al., 2017). Astrocytes express ten different isoforms of GFAP, which form the intermediate filament system with vimentin, nestin, and synemin (De Pablo et al., 2013; Hol & Pekny, 2015). Previous reports indicate that an increased number of GFAP+ astrocytes accompanies aging (Clarke et al., 2018; Cotrina & Nedergaard, 2002; Kohama et al., 1995; Nichols et al., 1993; J. D. Rodríguez-Callejas et al., 2023; Rozovsky et al., 1998; Y. Wu et al., 2005; Yoshida et al., 1996). Other reports show that astrocytes suffer a decrease in the length, volume, and branch points of their processes during aging, known as atrophy. Atrophic astrocytes have been described in aged non-human primates, humans, and mouse models of Alzheimer’s disease (AD) (Jyothi et al., 2015; Kanaan et al., 2010; Rodríguez et al., 2014; J. D. Rodríguez-Callejas et al., 2023). Until now, the exact causes that promote astrocytic atrophy are unknown.

Tree shrews (*Tupaia belangeri*) are small body-sized omnivorous mammals that belong to the *Scandentia* order (Fuchs, 2015). Tree shrews inhabit the tropical forests and plantation areas in Southeast Asia (Fuchs & Corbach-Söhle, 2010). They have been used as a biomedical model for multiple diseases such as hepatitis, hepatocellular carcinoma, myopia, psychosocial stress, depression, and breast cancer (Cao et al., 2003; Yao, 2017). Tree shrew is a promising species for the study of aging since they present a longer lifespan compared to rodents (-8 years) but shorter than most primates (Fuchs & Corbach-Söhle, 2010; Keuker et al., 2005). Also, they present a high homology with multiple human proteins, among which are proteins related to AD (i.e., Aβ, tau, APP) (Fan et al., 2018; H. Meyer et al., 2000; U. Meyer et al., 1998; Palchaudhuri et al., 1998, 1999; Pawlik et al., 1999). In old and aged tree shrews, there is an accumulation of iron in the brain parenchyma, higher RNA oxidation, and tau hyperphosphorylation in the hippocampus (Rodriguez-Callejas et al., 2020). Therefore, the tree shrew is a promising animal model to understand better the cellular processes related to brain aging. In the present study, we analyzed the density and morphology of GFAP+ astrocytes in the dorsal hippocampus of male tree shrews [dentate gyrus (DG), cornu ammonis 3 (CA3), CA2-CA1, and proximal subiculum (SUB)]. We also determined the rate of RNA oxidation by 8-hydroguanosine (8OHG), and the presence of S100A10, a marker of neuroprotective astrocytes. We observed that the number of GFAP+ astrocytes increased with age in most hippocampal regions, but it was significant only in CA3. In DG, CA3, and CA2-CA1, there was an increase in the number of astrocytes with atrophic features. SUB did not show significant changes in the astrocytic morphology across aging. During aging, we observed an enhanced number of GFAP+astrocytes with oxidized RNA in all regions analyzed, but only CA3 and CA2-CA1 regions showed a significant difference in aged animals compared to younger ones. S100A10, a calcium-binding protein associated with the neuroprotective role of astrocytes (Clarke et al., 2018), was present in the nucleus of astrocytes in adult animals. However, labeling of S100A10 in old and aged animals was rather cytoplasmic. Furthermore, aged tree shrews showed abundant atrophic GFAP+ astrocytes with cytoplasmic S100A10 labeling. Finally, cytoplasmic inclusions of S100A10 colocalized with Crm-1, a nuclear export protein. These results suggest that the transport of nuclear S100A10 in astrocytes could be associated with their morphological atrophy during the aging process in the tree shrew.

## 2. Methods

### 2.1 Subjects

Experimentally naive adult male tree shrews (*Tupaia belangeri*) were obtained from the breeding colony at the German Primate Center (Göttingen, Germany). Animals were housed individually under standard conditions complying with the European Union guidelines for the accommodation and care of animals used for experimental and other scientific purposes (2007/526/EC) on a 12 h light 12 h dark cycle with *ad libitum* access to food and water (Fuchs & Corbach-Söhle, 2010). All animal experiments were performed in accordance with the German Animal Welfare Act, which strictly adheres to the European Union guidelines (EU directive 2010/63/EU). Experienced veterinarians and caretakers constantly monitored the animals. The experiments were approved by the Lower Saxony State Office for Consumer Protection and Food Safety (LAVES, Oldenburg, Germany). Animals did not present neurological disorders or other injuries that could cause trauma to the central nervous system.

### 2.2 Tissue preparation

In the current study, we analyzed the brains of male tree shrews classified as adults (n = 2, mean age 3.8 years), old (n = 2, mean age six years), and aged (n = 2, mean age 7.5 years) according to previous literature (Fan et al., 2018; Keuker et al., 2004, 2005; Z. C. Wu et al., 2019). Animals were anesthetized with an i.p. injection (0.1 ml/100 g body weight) of GM II (ketamine, 50 mg/ml; xylazine 10mg/ml; atropin 0.1 mg/ml), and after the loss of consciousness, they received an i.p. injection of ketamine (400 mg/kg body weight). Bodies were transcardially perfused with cold (4 °C) saline (0.9 % NaCl) for 5 min. Subsequently, for fixation of the brains, cold (4 °C) 4 % paraformaldehyde (PFA) in 0.1 M phosphate buffer, pH 7.2, was infused for 15 min. The brains were removed and post-fixed in fresh 4 % PFA at 4 °C, where brains were stored until sectioning. Four days before sectioning, tissue was washed with 0.1 M phosphate-buffered saline (PBS: 0.15 M NaCl, 2.97 mM Na_2_HPO_4_-7H_2_O, 1.06 mM KH_2_PO_4_; pH 7.4) and immersed in 30 % sucrose in PBS at 4 °C. Horizontal sections (40 μm) were obtained from the dorsal hippocampal formation according to (Keuker et al., 2003) and series were prepared every 6th section (at intervals of 240 μm) using a sliding microtome (Leica RM2235). All brain sections were immediately immersed in cryoprotectant solutions for light microscopy [300 g sucrose (JT Baker); 400 mL 0.1M PB, and 300 mL ethylene glycol (Sigma) for 1 L] and for immunofluorescence [300 g sucrose; 10 g polyvinyl-pyrrolidone (PVP-40, Sigma); 500 mL of 0.1M PB and 300 mL ethylene glycol, for 1 L] and stored at -20°C until use in free-floating immunohistochemistry and immunofluorescence protocols.

### 2.3 Double and triple labeling immunofluorescence

To reduce autofluorescence and background due to PFA fixation, the dorsal hippocampus sections were incubated with 1% sodium borohydride (NaBH4) in PBS-1X for 10 min. Sections were rinsed with 0.5 % PBS-Tween20 twice for 3 min. To block potential nonspecific antibody binding the sections were incubated for 30 min with a solution containing 0.05 % Tween20, 0.1 % TritonX-100, 50 mM glycine, 2 % donkey serum, and 0.1 % BSA diluted in PBS-1X. Primary antibodies (supplementary table 1) were incubated in an antibody signal enhancer (ASE) solution (Flores-Maldonado et al., 2020; Rosas-Arellano et al., 2016), consisting of 0.05 % Tween20, 0.1 % TritonX-100, ten mM glycine, and 0.1 % hydrogen peroxide in PBS, and left overnight at 4 °C. The next day, sections were washed with 0.5 % PBS-Tween20 and incubated with secondary antibodies (see supplementary table 1) diluted in 0.1 % PBS-Tween20 for 2 h at RT. All sections were incubated with DAPI (1:1000, Invitrogen) in 0.2 % PBS-triton for 30 min. To reduce lipofuscin autofluorescence brain sections were incubated in 0.1% Sudan black for 15 min. Finally, sections were washed and mounted on glass slides with mounting medium VectaShield (Vector Laboratories).

### 2.4 Image acquisition

Images were obtained by a confocal microscope (Leica TCS-SP8) equipped with Diode (405 nm), OPSL (488 nm), OPSL (552 nm), and Diode (638 nm) laser. For Sholl analysis of GFAP+ astrocytes, a 63X objective was used. For double (GFAP versus 8OHG/S100A10) and triple labeling (GFAP versus S100A10 versus Crm-1) image acquisition, a 63X objective was used. All confocal images were obtained as z-stacks of single optical sections. Stacks of optical sections were superimposed as a single 2D image using the Leica LASX software. We captured images from different regions of the hippocampus (DG, CA3, CA2-CA1, and SUB) according to tree shrew neuroanatomical description (Keuker et al., 2003).

### 2.5 Sholl analysis

To quantify the length, volume, and number of branch points of astrocytic processes, we performed Sholl analysis using NeuronStudio software (Rodriguez et al., 2003). We captured five images per brain region (DG, CA3, CA2-CA1, and SUB) per subject for this analysis. A total of 20 images per slice were obtained. Ten astrocytes were analyzed for every image, giving a total of 1200 astrocytes to be analyzed. For GFAP versus S100A10 analysis, three images were captured per region (12 images per slice). All the images were captured with the following parameters: for GFAP optical gain 774, offset – 2.0 and 1.0 AU pinhole diameter; S100A10, optical gain 788, offset – 2.0 and 1.0 AU pinhole diameter. For Sholl analysis, image z-stacks were imported to NeuronStudio software, where astrocyte processes were reconstructed. After reconstruction, Sholl analysis was performed, which is based on the generation of concentric spheres radiating from the cell center, and at each sphere, the length, volume, and number of branch points of astrocytic processes are quantified. In our experiment, the radius of every concentric sphere is 1 µm larger than the previous. With the data of processes length in every concentric sphere, we generated the graphics of astrocytic processes length (y-axis) vs. radius (x-axis) (fig. 2A). To calculate the total astrocytic processes length (APL) (fig. 2A), total astrocytic processes volume (APV) and the number of astrocytic branch points (ABP) (fig. 2B and C, respectively) we sum the values of the respective parameter (length, volume or branch points) obtained in all the concentric spheres (radius) of the cell analyzed.

**Figure 1.**
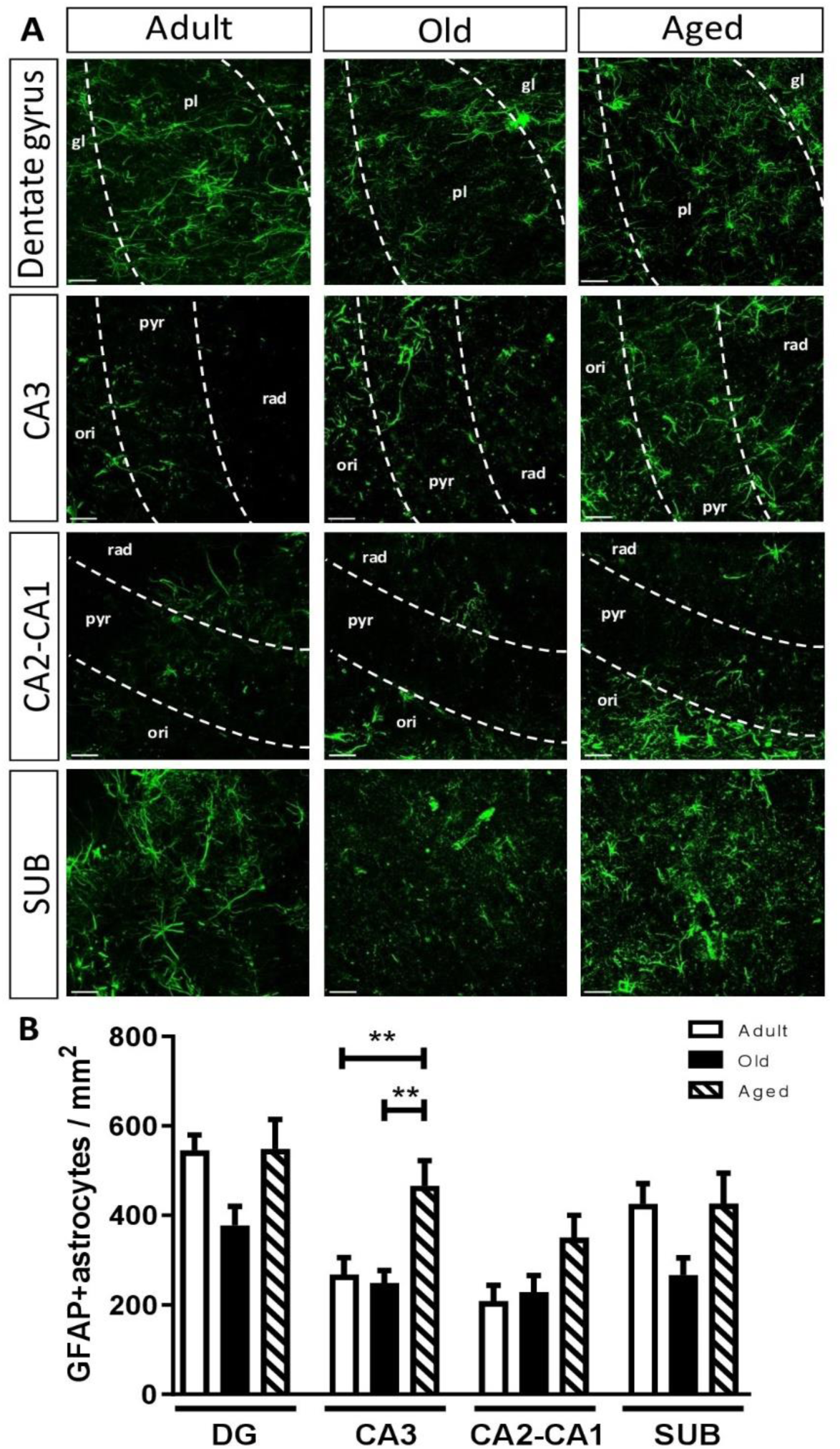
Quantification of GFAP+ astrocytes during aging in the hippocampus of three shrews. **(A) Representative images of GFAP+ astrocytes from DG, CA3, CA2-CA1, and proximal SUB of adult, old, and aged tree shrews**. The dotted lines delimit the subregions of DG (gl: granular layer; pl: polymorphic layer), CA3 (ori: *oriens stratum*; pyr: *pyramidale stratum*; rad: *radiatum stratum*), and CA2-CA1 (ori: *oriens stratum*; pyr: *pyramidale stratum*; rad: *radiatum stratum*). (B) The number of GFAP+ astrocytes was quantified in the dentate gyrus (DG), CA3, CA2-CA1 regions, and subiculum (SUB) of adult, old, and aged tree shrews. The number of GFAP+ astrocytes changes with aging, but it only reached a significant difference in CA3 regions (aged vs. adult and old). Data represents means ± SEM One-way ANOVA, Tukey post hoc analysis. *p < 0.05, **p < 0.01.

**Figure 2.**
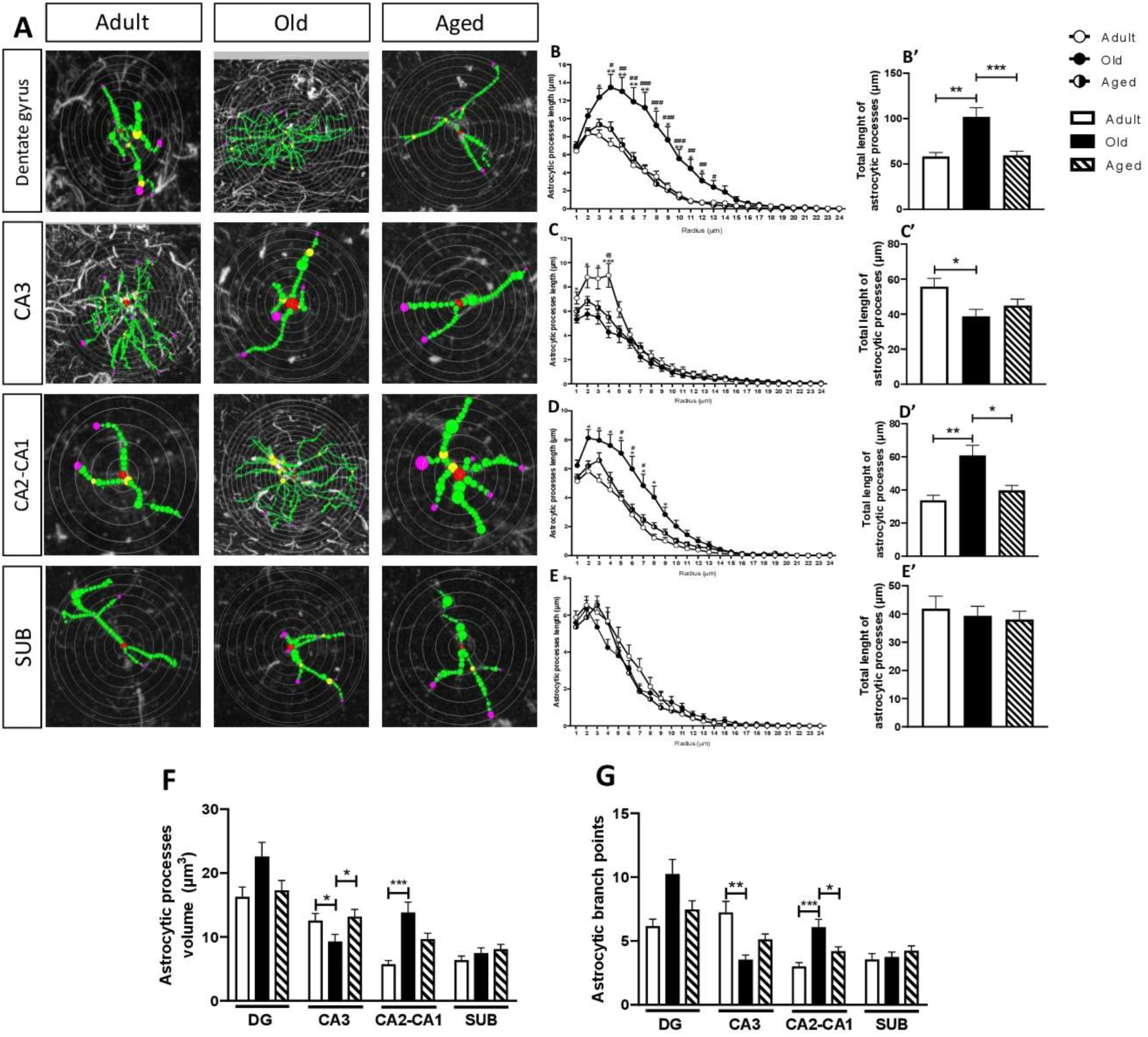
Morphological features of astrocytes in the hippocampus of tree shrews during aging. APL, APV, and ABP alterations increase in old tree shrews while decreasing in aged subjects. (**A)** Representative images of the Sholl method on GFAP+astrocytes of adult, old, and aged tree shrews. Note that in DG and CA2-CA1, astrocytes of old tree shrews present longer astrocytic processes than astrocytes of adult subjects. However, the astrocytes of aged subjects were smaller compared to old ones. In CA3, the size of astrocytes from old and aged subjects decreased compared to adults. The size of astrocytes from SUB did not change during the aging of tree shrews. Scale bar 5 µm. (**B-E)** Astrocytic processes length (APL) by Sholl curves. In DG (**B**), APL was significantly higher in old compared to adult (radius 3-12) and aged (radius 4-13) subjects. In CA3 (**C**), APL significantly decreased in old (radius 1-4) and aged (radius 4) subjects compared to adults. In CA2-CA1 (**D**), APL was significantly higher in old compared to adult (radius 2-9) and aged (radius 5-7) subjects. In SUB (**E**), no significant differences were detected. Data represents means ± SEM Multiple t-tests followed by Holm-Sidak post hoc analysis. * represent significant differences between old and adult groups. # represents significant differences between old and aged groups. @ represents differences between adult and aged groups. One symbol p < 0.05, two p < 0.01, and three p < 0.001. (**B’-E’)** Quantification of total APL. In GD (**B’**) and CA2-CA1 (**D’**), the total APL significantly increased compared to adults (GD: p < 0.01; CA2-CA1: p < 0.01), while decreased in aged subjects compared to old (GD: p < 0.001; CA2-CA1: p < 0.05). In CA3 (**C’**), the total APL of old subjects significantly decreased compared to adults (p < 0.05). Finally, the total APL did not change with aging in SUB (**E’**). (**F)** Astrocytic processes volume (APV) in all the regions analyzed in adult, old, and aged tree shrews. (**G)** Astrocytic processes branch points (ABP) in all the regions analyzed in adult, old, and aged tree shrews. Data represents means ± SEM One-way ANOVA followed by Tukey post hoc analysis. *p < 0.05, **p < 0.01, ***p < 0.001.

### 2.6 Number and percentage of 8OHG astrocytes per area

To determine if the number of GFAP+ astrocytes with RNA oxidation increases with age, we performed double labeling of GFAP versus 8OHG. Three images were captured on each region analyzed (DG, CA3, CA2-CA1, and SUB) at 63x. All the images were captured with the following parameters: GFAP (optical gain 774 and offset – 2.0) and 8OHG (optical gain 680, offset – 1.0). The counting of 8OHG+ astrocytes was performed using ImageJ software (NIH) Plugins--- Analyze---Cell counter). The sum of 8OHG+ astrocytes in a specific hippocampal region was divided by the total area analyzed. The total area analyzed was calculated by multiplying the area of a 63x image (0.061 mm^2^) by three (the number of images captured on every hippocampal region).

To calculate the percentage of 8OHG+ astrocytes, the sum of 8OHG+ astrocytes was multiplied by 100, and the product was divided by the total number of GFAP+ astrocytes in the region analyzed.

### 2.7 Classification and quantifications of S100A10+ / S100A10-astrocytes

To determine the proportion and morphological changes of S100A10+ and S100A10-astrocytes, a double-label immunofluorescence of GFAP versus S100A10 was performed. Three images were captured on each region analyzed (DG, CA3, CA2-CA1, and SUB) with a 63x objective. Two slices per subject were analyzed. All the images were captured following the parameters mentioned in section 2.5. For all the images, GFAP+ astrocytes were classified into four types: astrocytes with nuclear S100A10 labeling (nS100A10+), astrocytes with cytoplasmic S100A10 labeling (cS100A10+), astrocytes with nuclear and cytoplasmic S100A10 labeling (ncS100A10), and astrocytes without S100A10 labeling (S100A10-). After this classification, the total number of S100A10+/S100A10-astrocytes was performed using ImageJ software (Plugins---Analyze---Cell counter).

The percentage of S100A10+ / S100A10-astrocytes was calculated using the sum of S100A10+ / S100A10-astrocytes multiplied by 100 and the product was divided by the total amount of GFAP+ astrocytes in the region analyzed.

We performed a Sholl analysis of S100A10+ / S100A10- astrocytes and calculated APL, APV, and ABP as described in section 2.5.

### 2.8 Statistical analysis

Statistical analysis was performed using GraphPad Prism 9.0 software. The Shapiro-Wilk test was used to know if the data had a normal distribution. One-way ANOVA followed by a Tukey’s as posthoc test was performed, except for the quantification of total APL, APV, and ABP (Figs. 2A-C), where Kruskal-Wallis ANOVA test was followed by a Dunn’s test. For the Sholl analysis of APV, a multiple-t test followed by Holm-Sidak analysis was used. Differences were considered statistically significant when p ≤ 0.05. Data are presented as means ± SEM.

## 3. Results

### 3.1 Astroglial density during aging in the tree shrew

Astroglial density was evaluated in the four regions of the dorsal hippocampus (DG, CA3, CA2-CA1, and SUB) in the tree shrews at different ages (Fig. 1). We chose the dorsal hippocampus as this region has been related to cognitive functions (i.e., learning and memory) (Bannerman et al., 2004; Fanselow & Dong, 2010), but its alterations promote cognitive decline (Reichel et al., 2017). Quantification of GFAP+ astrocytes in DG, CA2, and SUB showed no significant differences with aging. However, in CA3, the number of GFAP+ astrocytes increased in aged subjects compared to adult and old tree shrews (Fig 1B). Despite no significant changes in the density of astrocytes during aging, we noted that the extension and morphology of the astrocytic processes were modified in old and aged tree shrews. Therefore, we performed a Sholl analysis in the regions of interest.

### 3.2 The length, volume, and branch points of astrocytic processes change with age in the hippocampal formation of the tree shrews

To analyze the morphological features of astrocytes during aging in the tree shrew, we performed a Sholl analysis of GFAP+ astrocytes in the four regions of interest. We quantified the length, volume (APV), and branch points (ABP) of astrocytic processes of adult, old, and aged tree shrews in the dentate gyrus (DG), CA3, CA2-CA1, and SUB. Representative images of GFAP+ astrocytes analyzed by the Sholl method are shown in Figure 2A.

Quantification of astrocytic processes length (APL) from the distance to soma showed a maximum between radius 2 and 4 in adults (white circles), old (black circles), and aged (white and black circles) tree shrews in all regions analyzed, declining thereafter until radius 24. In DG, astrocytes of old tree shrews had the largest APL in radius 4 to 12 compared to old animals, and in radius 3 to 12 compared to aged animals (Fig. 2B). In CA3, the APL was larger in adults in radius 1-4 compared to old, and in radius 4 compared to aged animals (Fig. 2C). In CA2-CA1, the largest APL was found in old subjects in radius 2-9 compared to adult and in radius 5-7 compared t aged animals (Fig. 2D). In SUB, APL had a similar complexity among all age groups (Fig. 2E).

Then, we quantified the total APL in all the regions of interest. In DG, the largest total APL was seen in astrocytes from old animals, compared to adult and aged ones, indicating hypertrophy in old animals, followed by atrophy in aged subjects (Fig. 2B’). In CA3, total APL was highest in adult animals, decreasing in old and aged subjects (Fig. 2C’). In CA2-CA1, total APL was the highest in old animals, decreasing in aged subjects (Fig. 2D’). In SUB, there were no differences in total APL among the three age groups, and APL was relatively low (40 µm) (Fig. 2E’).

We also quantified the astrocytic process volume (APV). In DG and SUB, no significant differences in APV existed among age groups. In CA3, APV decreased in old animals compared to adults and aged animals. Conversely, in CA2-CA1, a larger APV was observed in old animals compared to adults and aged animals (Fig. 2F).

The other morphological parameter analyzed was the astrocytic branch points (ABP). No significant differences were seen in DG and SUB among age groups. In CA3, astrocytes from old subjects showed a smaller APV compared to adults. In CA2-CA1, astrocytes of old subjects had the highest APV than the other two age groups (Fig. 2G).

These results indicate that astrocytes present morphological changes that are regional dependent during the aging process in the tree shrew. In DG and CA2-CA1, hypertrophy is followed by atrophy, while in CA3, atrophy is present in old animals. Previous studies suggest that age-related oxidative stress could promote astroglial atrophy. Thus, we analyzed the levels of RNA oxidation in astrocytes of the hippocampal region.

### 3.3 Increased RNA oxidation in astrocytes during aging in the three shrew

8OHG is produced by oxidation of the RNA, and it is considered an early marker of oxidative stress in the brain. To determine the role of RNA-oxidation in the atrophy of astrocytes, we conducted double-label immunofluorescence of GFAP and 8OHG (Fig. 3). GFAP+ astrocytes of adult tree shrews do not present 8OHG label (Fig. 3A top row, white arrows). However, in old (Fig. 3A middle row) and aged (Fig. 3A bottom row) animals, most GFAP+ astrocytes present cytoplasmic 8OHG label (white arrowheads), mainly located around the nucleus. We quantified the number of GFAP+ / 8OHG+ astrocytes in all regions analyzed. The number of GFAP+ / 8OHG+ astrocytes was higher in aged subjects compared to younger animals in all regions analyzed, being significantly different in CA3 and CA2-CA1 (Fig. 3B). As density of GFAP+ astrocytes was highest in aged animals (mainly in CA3, see Fig. 1), we calculated the percentage of GFAP+/8OHG+ astrocytes in all the regions. Notably, most astrocytes in old tree shrews showed substantial RNA oxidation, with a percentage close to 80 % in all regions analyzed. The percentage remained high in aged subjects, showing a significant difference in CA2-CA1 (adult vs. old and adult vs. aged) and SUB (adult vs. aged) (Fig. 3C).

**Figure 3.**
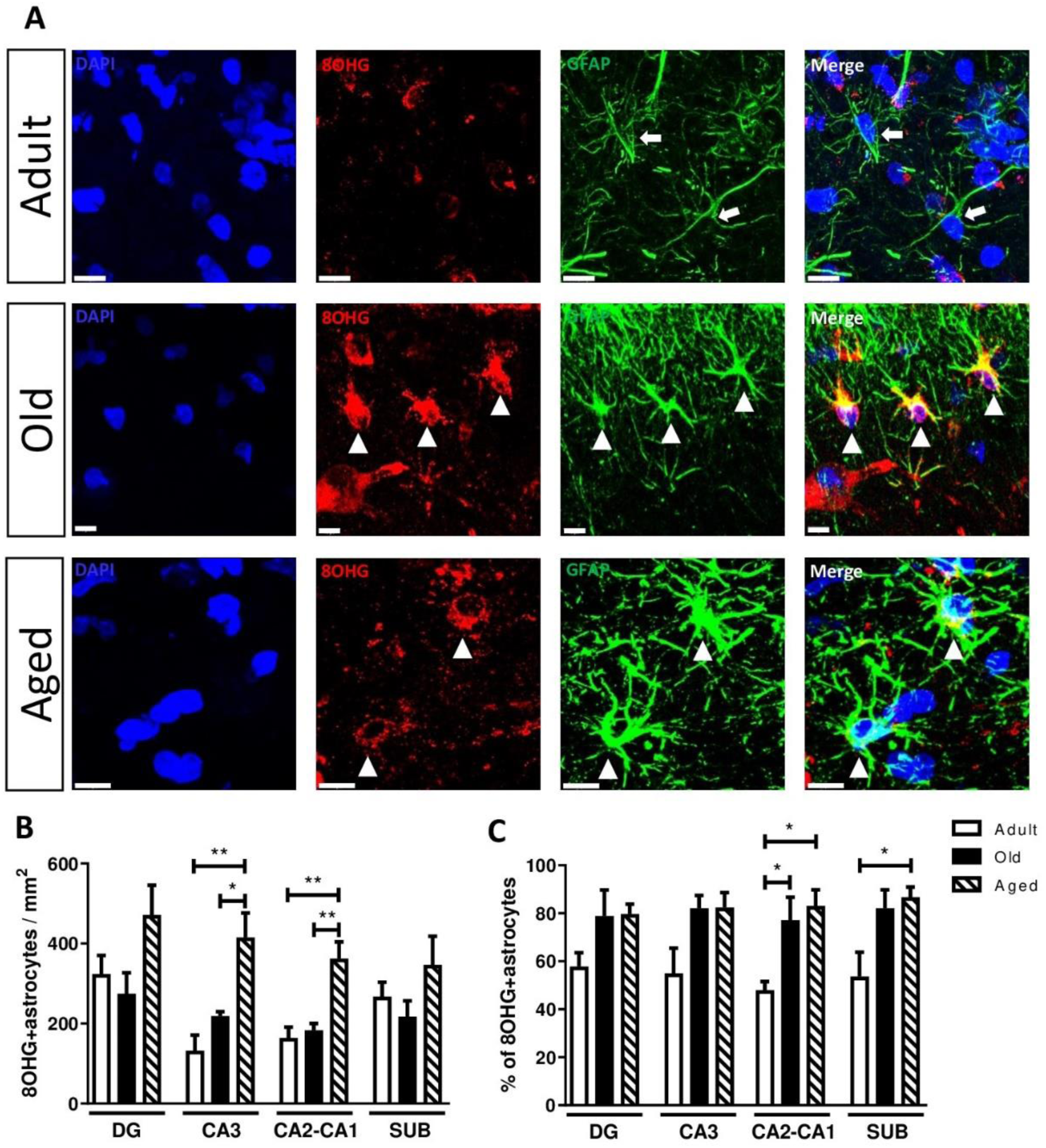
Oxidized-RNA in GFAP+astrocytes in the hippocampus of tree shrews. **(A)** Representative images of GFAP+ astrocytes and 8OHG in the hippocampus of adult, old, and aged tree shrews. In adult tree shrews, a few GFAP+ astrocytes (green) showed 8OHG-labelling (red) (marked with white arrows). However, the number of GFAP+ astrocytes/8OHG+ increased in old and aged animals (marked with white arrowheads). DAPI was used as a nuclear counterstain. Scale bar 10 μm. (**B)** Quantification of GFAP+astrocytes with RNA oxidation. The total number of astrocytes with oxidized RNA increased with age in all the regions analyzed; however, it reached significant differences only in CA3 and CA2-CA1. (**C)** The percentage of GFAP+astrocytes with RNA oxidation. In the four regions analyzed, the percentage of astrocytes with oxidized RNA increased in old and aged tree shrews compared to adults, but it was significant only in CA2-CA1 and SUB. Data represents means ± SEM. One-way ANOVA, Tukey post hoc analysis. *p < 0.05, **p < 0.01.

### 3.4 S100A10 expression in GFAP+ differs during aging in three shrews

Previous studies demonstrate that S100A10 is a marker of a neuroprotective astrocytic activation (Clarke et al., 2018). Moreover, astrocytes lacking S100A10 had an atrophic morphology (J. D. Rodríguez-Callejas et al., 2023). We used double labeling (S100A10 and GFAP) to determine if the atrophic phenotype of GFAP+ astrocytes is associated with altered S100A10 expression.

We noticed three types of S1000A10 labeling in GFAP+astrocytes (Fig. 4): nuclear S100A10 (nS100A10+) (Fig. 4 top row); cytoplasmic S100A10 (cS100A10+) (Fig. 4 second row); and nuclear/cytoplasmic S100A10 (ncS100A10) (Fig. 4 third row). In addition, there were GFAP+ astrocytes that lack the S100A10 label (S100A10-) (Fig. 4 bottom row). When comparing the morphological features of those four types of S100A10-labeled astrocytes, we found regional differences in the percentage of S100A10+/S100A10- astrocytes with aging (Figure 5 – 8, panels B). In adult and old animals, the type of astrocytes that predominates were nS100A10+ followed by S100A10- in all the regions analyzed. However, in aged tree shrews, the percentage of nS100A10 decreased, while the percentage of cS100A10+ and ncS100A10+ types increased. Thus, the number of astrocytes with cS100A10 label increased in aged tree shrews.

**Figure 4.**
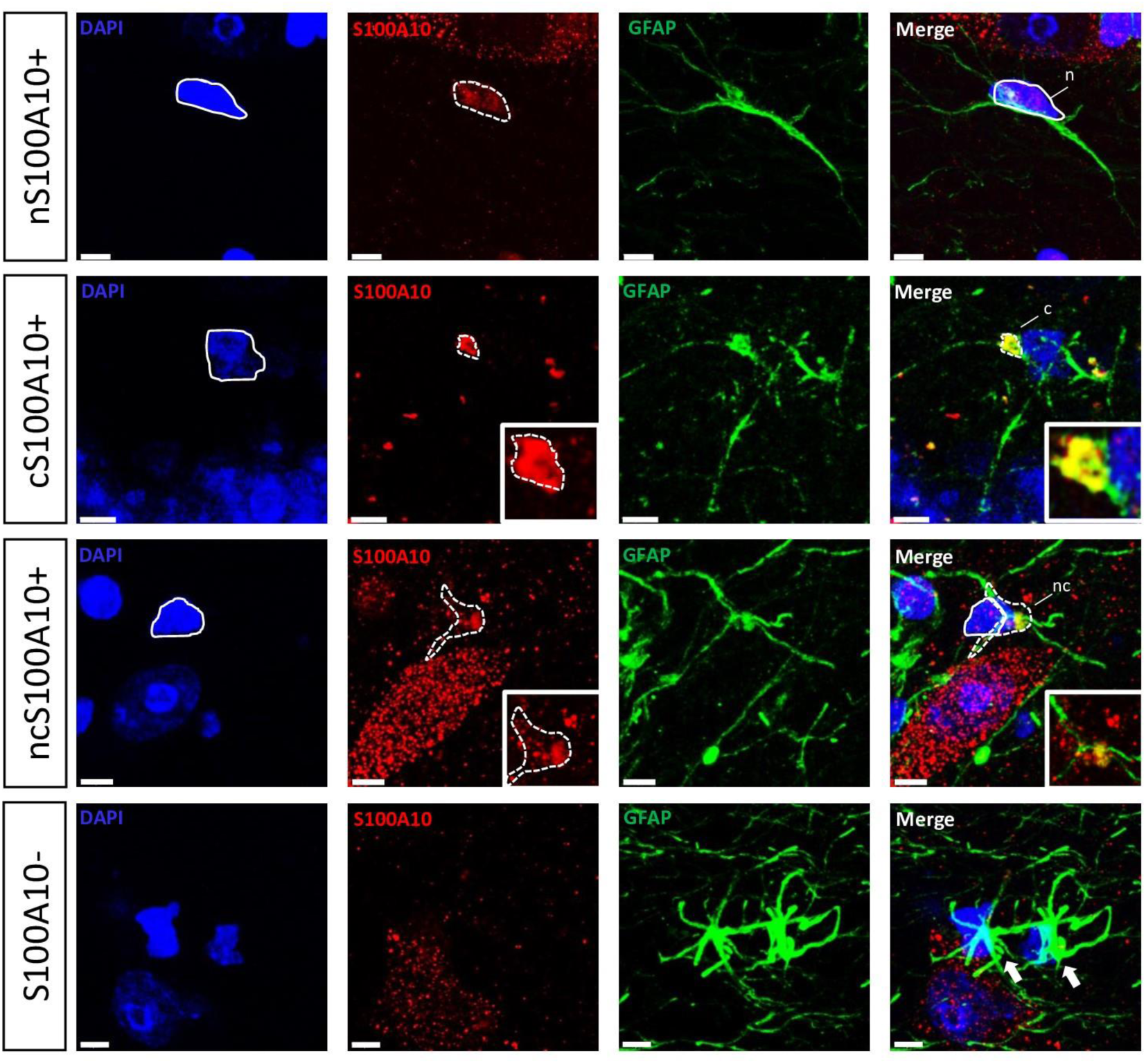
S100A10+ and S100A10- astrocytes in the hippocampus of adult and aged tree shrews. Double label immunofluorescence of GFAP (green) and S100A10 (red) in the hippocampus of tree shrews. S100A10 showed four types of labeling in astrocytes: nuclear label (nS100A10+); cytoplasmic label (cS100A10+); nuclear and cytoplasm label (ncS100A10+); and lack of label (S100A10-; white arrow). Squares indicate high magnifications of middle panels. DAPI was used as a nuclear counterstain. Scale bar 5 μm.

**Figure 5.**
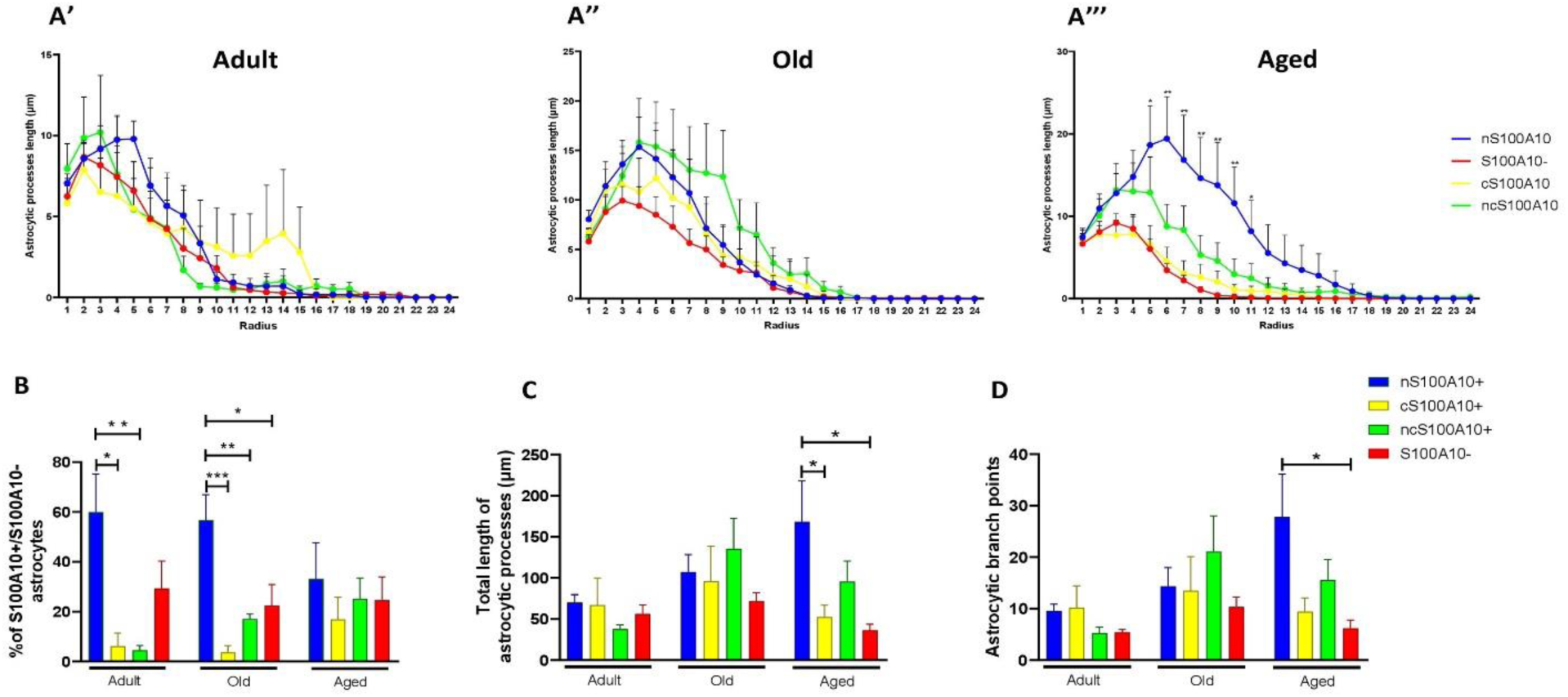
Sholl analysis of S100A10+ and S100A10- astrocytes of the DG. **(A)** Radial analysis of S100A10+/S100A10- astrocytic length in adult (A’), old (A"), and aged (A’") tree shrews. In old and aged tree shrews, APL of nS100A10+ and ncS100A10+ astrocytes tends to be higher than cS100A10+ and S100A10- astrocytes. Data represents means ± SEM Multiple t-tests followed by Holm-Sidak post hoc analysis. a: p < 0.05; b: p < 0.01. (**B)** Percentage of nS100A10+, cS100A10+, ncS100A10, and S100A10- astrocytes per area. In adult and old subjects, the percentage of nS100A10+ predominates over the other types of labels. However, in aged subjects, all types of astrocytes present similar percentages. (**C)** Total APL quantification. (**D)** ABP quantification. In adult and old tree shrews, significant differences in total APL and ABP were not detected between the types of astrocytes. In aged subjects, total APL and ABP of nS100A10+ astrocytes significantly increase compared to cS100A10+ and S100A10- astrocytes. Data represents means ± SEM One-way ANOVA, Tukey post hoc analysis. *p < 0.05, **p < 0.01.

A Sholl analysis was performed to determine if this change in the location of S100A10 in aged subjects is related to atrophy. A radial analysis of GFAP+ astrocytes from DG (Figure 5), CA3 (Figure 6), CA2-CA1 (Figure. 7), and SUB (Figure 8) in adult (A’), old (A’’) and aged (A’’’) tree shrews was done. The APL curves of the four types of S100A10 label in GFAP+ astrocytes in adult subjects did not show differences, which denotes that the size of the four astrocytic types was similar. In old subjects, the Sholl curve of nS100A10+ showed higher APL than the other three types of labels in CA3, CA2-CA1, and SUB (mainly in the radius 3-11). In the case of DG, the four Sholl curves present similar values. Finally, in aged tree shrews, DG and CA3 showed the highest APL in nS100A10+ astrocytes. In CA2-CA1, APL levels tend to be higher in nS100A10+ than the other three types of labeled astrocytes. In SUB, the highest levels of APL were seen in ncS100A10+. The quantification of total APL (Figures 5 - 8, panels C) corroborates the Sholl curves observed in the radial analysis.

**Figure 6.**
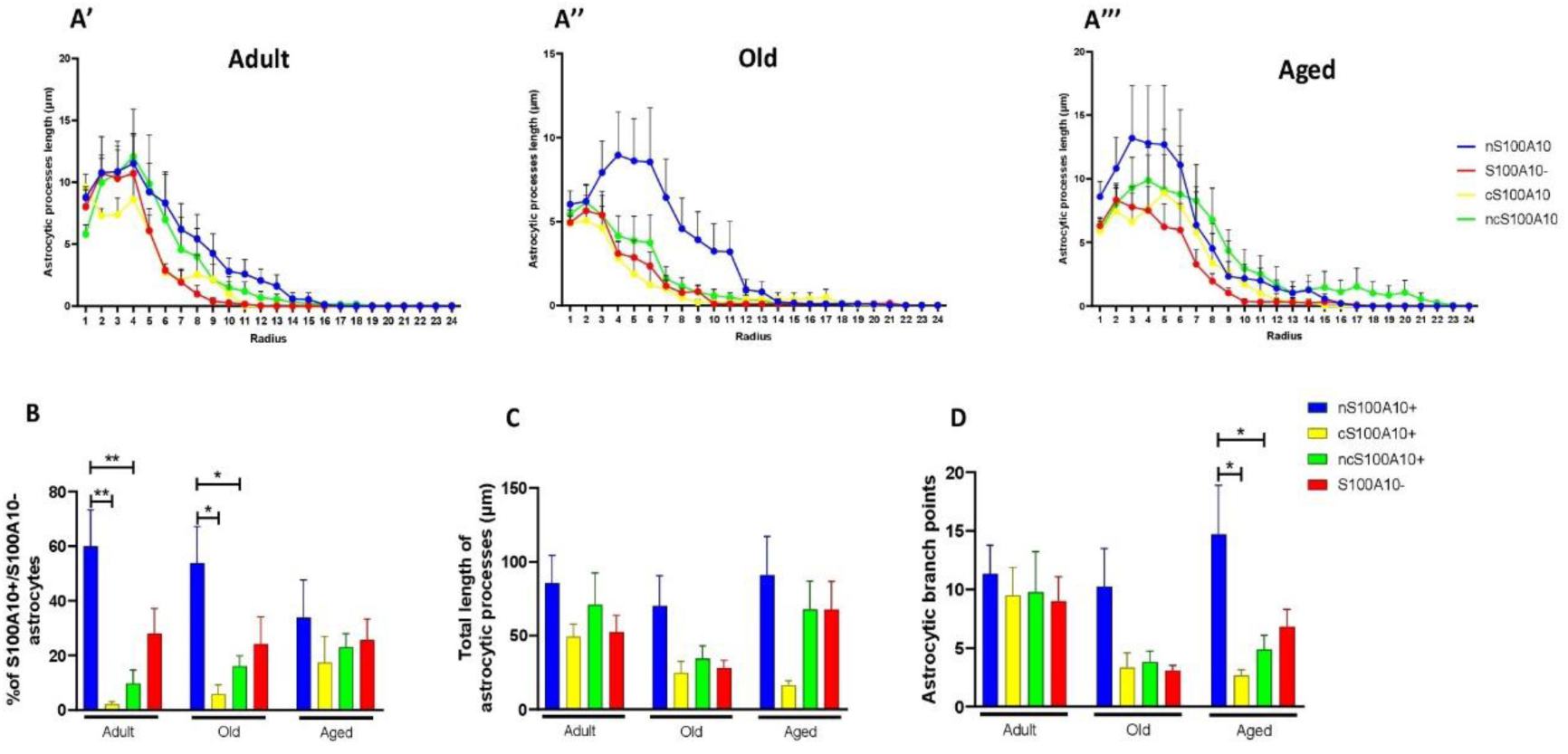
Sholl analysis of S100A10+ and S100A10- astrocytes of CA3. **(A)** Radial analysis of S100A10+/S100A10- astrocytic length in adult (**A’**), old (**A**"), and aged (**A’**") tree shrews. Data represents means ± SEM Multiple t-tests followed by Holm-Sidak post hoc analysis. (**B)** Percentage of nS100A10+, cS100A10+, ncS100A10, and S100A10- astrocytes per area. As observed in DG, nS100A10+ astrocytes predominate in adult and old subjects. Meanwhile, in aged subjects the four types of astrocytes present similar percentages. (**C)** Total APL quantification. In all the age groups, total APL tends to be higher in nS100A10+, ncS100A10, and S100A10- compared to cS100A10+ astrocytes. (**D)** ABP quantification. There were no differences in ABP in the four type of astrocytes in adult subjects. In old and aged subjects, the ABP of nS100A10+ increased compared to the other three types of astrocytes (significantly increases in aged subjects: nS100A10+ vs. cS100A10+ and ncS100A10+). Data represents means ± SEM One-way ANOVA, Tukey post hoc analysis. *p < 0.05, **p < 0.01.

**Figure 7.**
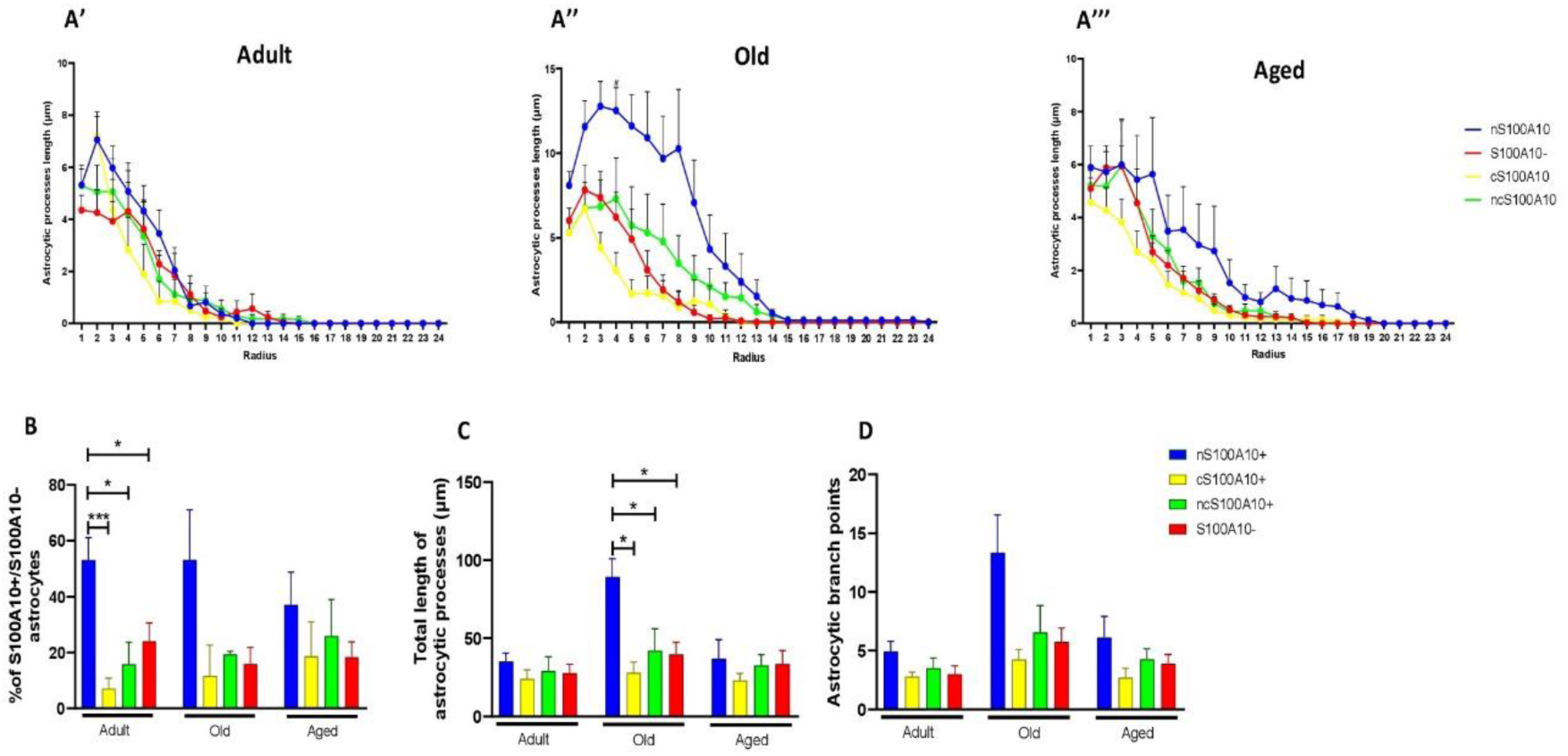
Sholl analysis of S100A10+ and S100A10- astrocytes of CA2-CA1 of adult, old, and aged tree shrews. **(A)** Radial analysis of S100A10+/S100A10- astrocytic length in adult (**A’**), old (**A**"), and aged (**A’**") tree shrews. In adult subjects, all the astrocytic types present similar levels of APL. While in old and aged tree shrews, the APL of nS100A10+ tends to be higher than the other types. Data represents means ± SEM Multiple t-tests followed by Holm-Sidak post hoc analysis. #: p < 0.05 between nS100A10+ group and cS100A10+ group. (**B)** Quantification of the percentage of nS100A10+, cS100A10+, ncS100A10, and S100A10- astrocytes per area. In adult and old subjects, the percentage of nS100A10+ astrocytes tends to be higher than the other types (with significant differences in adult subjects). In aged subjects, all types of astrocytes present similar percentages. (**C)** Total APL quantification. (**D)** ABP quantification. In adult and aged tree shrews, total APL and ABP significant differences were not detected between the four types of astrocytes. In old subjects, the total APL and ABP of nS100A10+ astrocytes were significantly higher than the other three types of astrocytes. Data represents means ± SEM One- way ANOVA, Tukey post hoc analysis. *p < 0.05, **p < 0.01, ***p < 0.001.

**Figure 8.**
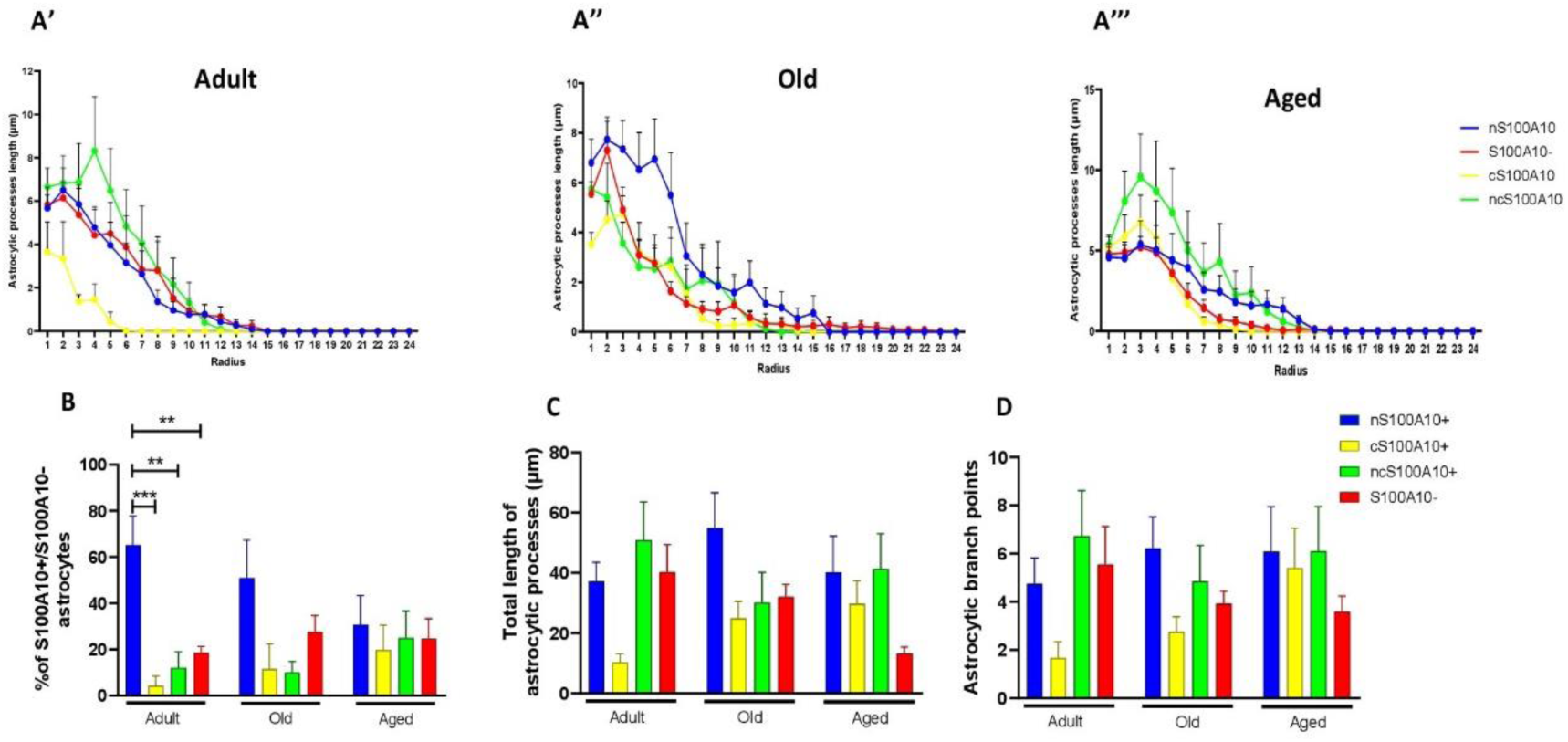
Sholl analysis of S100A10+ and S100A10- astrocytes of SUB of adult, old and aged tree shrews. **(A)** Radial analysis of S100A10+/S100A10- astrocytic length in adult (**A’**), old (**A**”), and aged (**A’**”) tree shrews. In old subjects, nS100A10+ astrocytes showed higher levels of APL than the other groups. In aged subjects, ncS100A10+ astrocytes present higher APL than the other types. Data represents means ± SEM Multiple t-tests followed by Holm-Sidak post hoc analysis. (**B)** Quantification of the percentage of nS100A10+, cS100A10+, ncS100A10, and S100A10- astrocytes per area. In adult and old subjects, the percentage of nS100A10+ astrocytes tends to be higher than the other types (with significant differences in adult subjects). In aged subjects, the four types of astrocytes present similar percentages. (**C)** Total APL quantification. (**D)** ABP quantification. In adults, cS100A10+ astrocytes present lower total APL and ABP than other astrocytes. In old subjects, total APL and ABP of nS100A10+ astrocytes tend to be higher than the other groups. In aged tree shrews, the morphological parameters of S100A10- astrocytes were lower compared to the other three types. Data represents means ± SEM One-way ANOVA, Tukey post hoc analysis. *p < 0.05, **p < 0.01, ***p < 0.001.

The quantifications of APV (Supplementary Figure 1) and ABP (Figures 5 – 8, panels D) present similar values in DG in adult and old subjects. However, in aged subjects, nS100A10+ showed the highest levels of APV and ABP (significant differences in ABP: nS100A10+ vs. cS100A10+ and nS100A10+ vs. S100A10-; significant differences of APV: nS100A10+ vs. S100A10-) than younger animals. Adult subjects had similar levels of ABP and APV in CA3, CA2-CA1, and SUB. However, in old and aged tree shrews, APV and ABP were higher in nS100A10+ astrocytes than the other three types of labels (ABP: nS100A10+ vs. S100A10- in old subjects; nS100A10+ vs. S100A10- in aged subjects; APV: nS100A10+ vs. cS100A10+ and nS100A10 vs. ncS100A10+ of aged subjects).

These results demonstrate that adult and old tree shrews had a predominance of nuclear S100A10 label astrocytes in the hippocampal formation, while aged subjects showed no differences between the four types of S100A10 label. Sholl analysis revealed that the size of nS100A10+ astrocytes was higher than the size of ncS100A10+ and cS100A10+ astrocytes in old and aged tree shrews, which suggests that the loss of S100A10 in the nucleus could be a factor that promotes the astroglial atrophy.

### 3.5 Cytoplasmic S100A10 colocalized with Crm-1 in astrocytes of aged three shrews

To know if the Crm-1 protein mediates the transport of S100A10 to the cytoplasm in atrophic astrocytes, we performed a triple-label immunofluorescence (GFAP/S100A10 / Crm-1; Figure 9). In adult tree shrews most astrocytes present Crm-1 label in the nucleus (Figure 9 top row). However, old and aged animals present cytoplasmic label of S100A10 that colocalized with Crm-1, close to the nucleus (Figure 9, middle and bottom rows). These results suggest that Crm-1, an export protein, could promote the age-dependent exportation of nuclear S100A10 to the cytoplasm in astrocytes of the hippocampal formation of the tree shrews.

**Figure 9.**
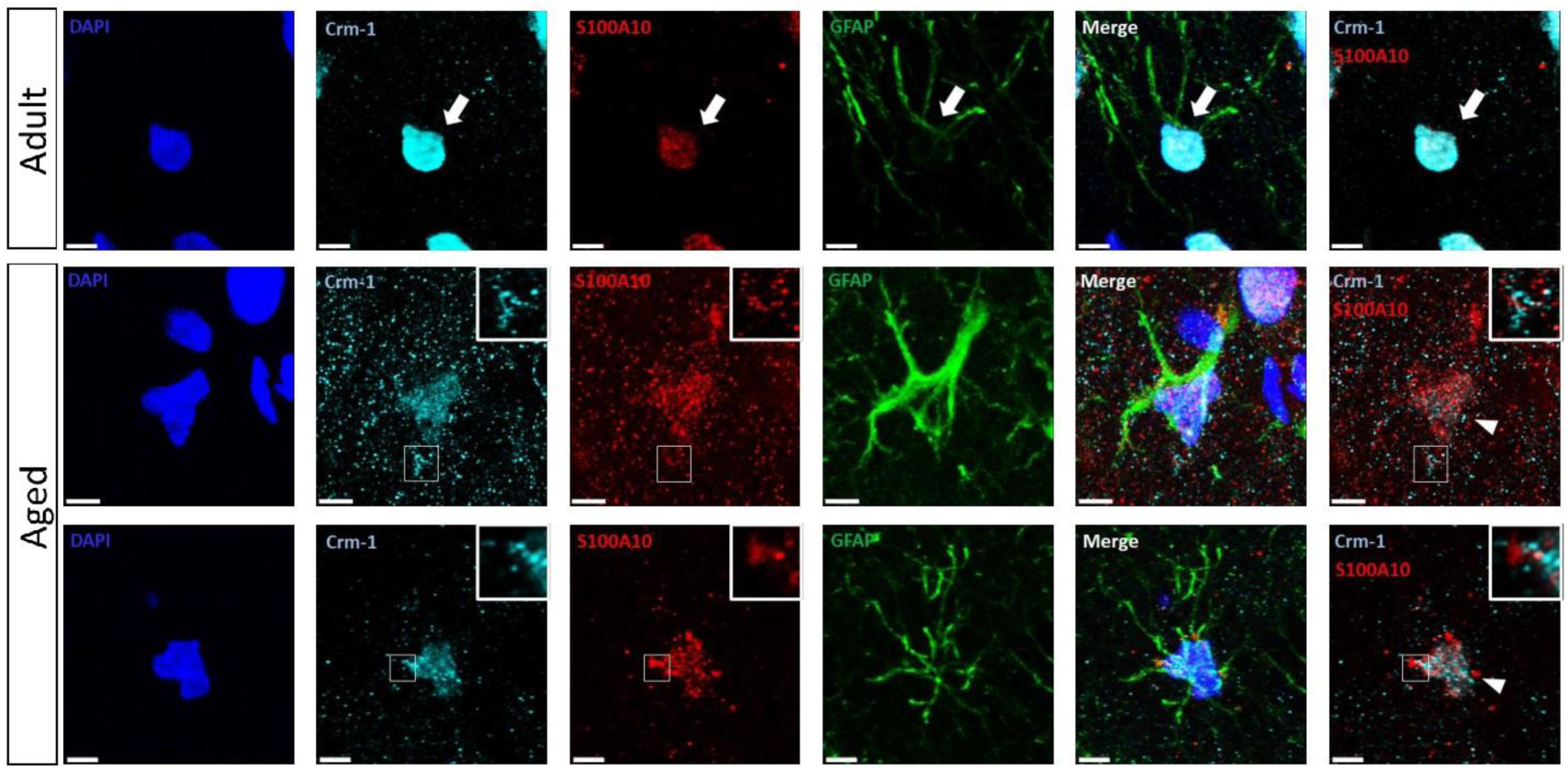
Nuclear and cytoplasmic S100A10 labeling in astrocytes. Triple label immunofluorescence for GFAP (green), S100A10 (red), and Crm-1 (cyan) in the hippocampus of adult and aged tree shrews. Most astrocytes of adult animals present positive label of S100A10 and Crm-1 in their nucleus (white arrow). In aged tree shrews, extra-nuclear S100A10 colocalized with Crm-1 label in the cytoplasm (middle and bottom panels). DAPI was used as a nuclear counterstain. Scale bar 5 μm.

**Figure 10.**
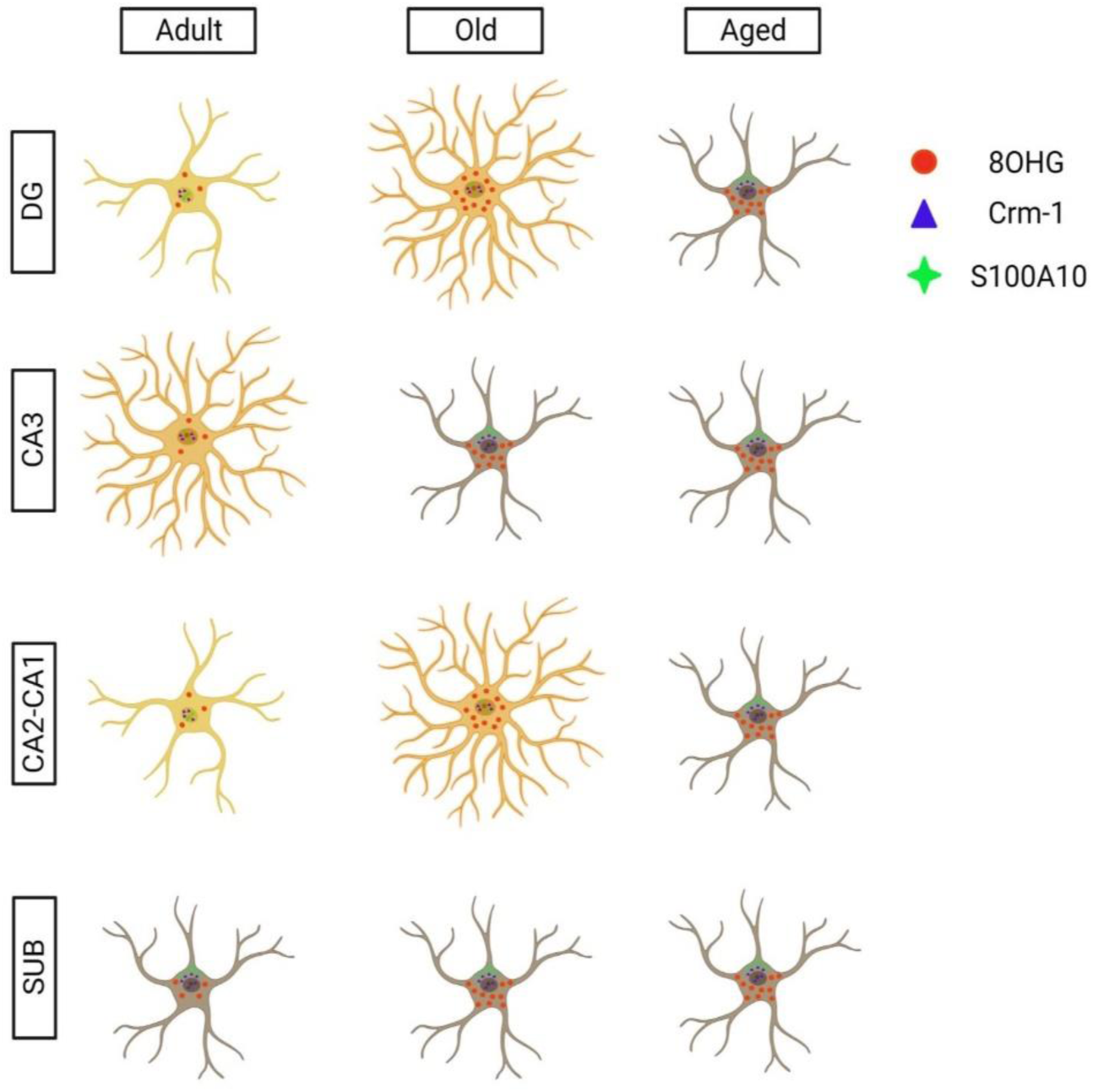
Age related alterations in hippocampal astrocytes of tree shrews. Astroglial activation in old tree shrews was followed by an increase in atrophic astrocytes in aged animals in DG and CA2-CA1 subregions of the hippocampus. In CA3 of adult animals, the number of hypertrophic astrocytes increased, followed by atrophy in old and aged subjects. However, astrocytes in the SUB did not show morphological differences during aging in the tree shrew. RNA oxidation (8OHG+) was present in astrocytes of adult, old, and aged animals in all the regions analyzed. S100A10 was located in the nucleus of astrocytes of adult and old animals in DG, CA3, and CA2-CA1. However, in aged subjects, S100A10 appears in the cytoplasmic compartment with Crm-1, mainly in atrophic astrocytes. This data may indicate that the export of proteins from the nucleus to the cytoplasm may be associated with the atrophy of the cells. Created with BioRender.

## 4. Discussion

### 4.1 Hypertrophic and atrophic astrocytes in the tree shrew

The total number of GFAP+ astrocytes did not change across aging in the hippocampal sub-regions of the three shrews, except in CA3, where aged animals had an enhanced number of astrocytes than younger subjects. However, several morphological alterations during the aging process differ between regions. In DG and CA2-CA1, there was a marked hypertrophic phenotype in old three shrews compared to adults; however, aged animals mainly presented astrocytes with shorter processes, resembling an atrophic phenotype. In CA3, old animals showed atrophic astrocytes that remain short in the aged group. Notably, in SUB, the morphological parameters remain unchanged along aging, with short processes already in adult animals. These results indicate that astroglial activation varies depending on the hippocampal sub-regions during aging in the tree shrews.

Hypertrophic astrocytes in old tree shrews could be promoted by age-related alterations occurring in those sub-regions, such as iron accumulation (Rodriguez-Callejas et al., 2020), oxidative stress, tau hyperphosphorylation, and Aβ plaques accumulation (Rodriguez-Callejas et al., 2020; Yamashita et al., 2010). In particular, SUB is a region that showed high levels of RNA oxidation, nuclear tau hyperphosphorylation, and iron accumulation, even since adulthood. Moreover, SUB is the region with the lowest levels of resting and activated iron-storage microglia (ferritin+ microglia) (Rodriguez-Callejas et al., 2020). Here, we showed that SUB was the region with a null astrocytic activation. SUB communicates to the anterior, lateral dorsal, and midline thalamic nuclei, retrosplenial cortex, and mammillary bodies (Aggleton & Christiansen, 2015), that renders SUB a fundamental region for visceral and emotional control. SUB’s lowest levels of glial activation (microglia and astrocytes) may leave this region vulnerable to age-related alterations in those functions. On the other hand, CA3 showed the largest astrocytes in adult animals, but in old and aged animals, there was a marked reduction in the morphological parameters, such as APL, APV and ABP, which denotes atrophy as previously observed (Rodríguez et al., 2014). Our previous study showed that CA3 presented the lowest levels of oxidized RNA, nuclear tau hyperphosphorylation, and iron accumulation compared to DG, CA2-CA1, and SUB. In addition, CA3 showed high levels of resting and activated ferritin+ microglia. Thus, the abundance of activated microglia and GFAP+ astrocytes in CA3 of aged tree shrews may be a compensatory mechanism against the astroglial atrophy. In the case of microglia, compensatory proliferation has been detected in mice models of facial nerve axotomy, as microglia proliferate for a longer time in old mice than young mice as a compensatory mechanism for the increased amount of dystrophic microglia (Conde & Streit, 2006).

The importance of regional differences in glial proliferation and activation has been observed in CA3 and CA1 regions during ischemia/hypoxia. CA3 and CA1 are contiguous regions interconnected by Schaffer collaterals (Andersen et al., 2007); However, in ischemia/hypoxia patients (Bartsch et al., 2010, 2015; Petito et al., 1987; Zola-Morgan et al., 1986) and animal models (Kirino, 1982; Pulsinelli et al., 1982) the CA1 neurons are more vulnerable to ischemia/hypoxia-related damage than CA3 neurons. Furthermore, previous studies in animal models demonstrate that in CA3 the number of astrocytes and microglia was higher than in CA1, which facilitates the phagocytosis of apoptotic pyramidal neurons and reduces the ischemia/hypoxia-related damages in CA3 compared to CA1 (Lana et al., 2014, 2017).

### 4.2 Enhanced RNA oxidation in GFAP+ astrocytes during aging in the tree shrew

Until now, the mechanism triggering astroglial atrophy has been unknown. In murine models, astrocytes of old subjects had a proinflammatory phenotype denoted by the overexpression of genes related to antigen processing and antigen presentation, complement cascade, defense response, metal ion binding, zinc ion binding, and cytokines (Boisvert et al., 2018; Orre et al., 2014). The enhanced number of astrocytes with a proinflammatory phenotype in aged animals and the increased number of activated microglia with a proinflammatory phenotype (Norden & Godbout, 2013; von Bernhardi et al., 2015) are related to a pro-oxidative environment promoting oxidative stress. An early alteration induced by oxidative stress is oxidation to the RNA, widely reported in old humans, rats, SOD1^G93A^ mice, marmosets, and tree shrews (Chang et al., 2008; Fiala et al., 1989; Nunomura et al., 1999; J. Rodríguez-Callejas et al., 2019; Rodriguez-Callejas et al., 2020). The most reliable marker of RNA oxidation is 8-hydroxyguanosine (8OHG) (Kasai et al., 2008; Syslová et al., 2014). In marmosets, tree shrews, and humans, the number of 8OHG positive cells (8OHG+) increases with age in the hippocampus, entorhinal cortex, temporal lobe, and parietal lobe (Ding et al., 2005, 2006; Nunomura et al., 1999, 2001, 2002, 2012; J. Rodríguez-Callejas et al., 2019; Rodriguez-Callejas et al., 2020). Also, in astrocytes of old rats, there is an increase in RNA oxidation compared to young rats (Bellaver et al., 2017). Thus, RNA oxidation accompanies the aging process. However, astrocytic atrophy and RNA oxidation do not entirely correlate in our study, since the appearance of atrophy was region-specific. In contrast, RNA oxidation increased with age in all the brain regions analyzed, which suggests that in tree shrews, RNA oxidation is not the cause of astrocytic atrophy during the aging process.

### 4.3 Loss of nuclear S100A10 promotes astroglial atrophy in aged three shrews

S100 is a family of 21 calcium sensor proteins encoded by individual genes found exclusively in vertebrates (Zimmer et al., 2013). S100A10 is expressed in most tissues but is highly expressed in the kidney, intestine, and lung (Saiki & Horii, 2019). Also, S100A10 expression has been detected in macrophages, fibroblasts, endothelial cells, epithelium, and cancer cell lines (Huang et al., 2003; Madureira et al., 2012; Phipps et al., 2011; Surette et al., 2011; Zokas & Glenney Jr, 1987). In the brain, S100A10 has been detected in the hippocampus, raphe nuclei, hypothalamus, and cerebral cortex, and it can bind to multiple ion channels and 5-HT_1B_ receptors (Svenningsson & Greengard, 2007).

Previous studies using models of ischemia have shown that reactive astrocytes promote tissue repair and recovery (Zamanian et al., 2012). This neuroprotective-type astrocytes overexpress S100A10 protein (Clarke et al., 2018). In the common marmoset, S100A10- astrocytes had an atrophic phenotype characterized by lower APL, APV, and ABP than S100A10+ astrocytes (J. D. Rodríguez-Callejas et al., 2023). Here, we observed four types of labeling in astrocytes: S100A10 label in the nucleus (nS100A10+), S100A10 label in the cytoplasm (cS100A10+), S100A10 label in the nucleus and cytoplasm (ncS100A10). and astrocytes negative to S100A10 label (S100A10-). nS100A10+ astrocytes predominate in adult subjects of all the regions analyzed. However, in old tree shrews and, to a greater extent, in aged subjects, the percentage of cS100A10+, ncS100A10+, and S100A10- astrocytes increased, which suggests that during aging, the transport of S100A10 outside the nucleus is increased. Subsequently, a Sholl analysis of the four types of astrocytes demonstrates that in old and aged tree shrews, the morphological parameters analyzed (APL, APV, and ABP) are lower in cS100A10+ and ncS100A10+ astrocytes compared to nS100A10+ astrocytes. This suggests that atrophic astrocytes lose the nuclear S100A10. Studies in tumor giant cells demonstrate that the SUMOylation of S100A10 promotes its transport to the nucleus, where it promotes the expression of genes such as *DEFA3*, *ARHGEF18*, and *PTPRN2*, which are related to cytoskeleton remodeling and actin dynamics (Zhao et al., 2021). However, the inhibition of SUMOylation promotes the loss of S100A10 in the nucleus, which reduces the proliferation and migration of these cells (Zhao et al., 2021). Thus, the loss of S100A10 in the nucleus of astrocytes in aged three shrews could alter the astroglial activation (loss of proliferation and migration) and could be associated with the loss of proteins related to cytoskeletal dynamics leading to the atrophy of the cell. Future studies should further characterize the relationship between S100A10 and astroglial activation or astrocytic atrophy.

### 4.4 Nuclear export protein Crm-1 colocalizes with cytoplasmic S100A10 in atrophic astrocytes

The chromosome region maintenance 1 protein (Crm-1) or exportin-1 (XPO-1) can bind to peptides of 8-15 residues known as nuclear export signals (NES) of multiple proteins to transport those cargos from the nucleus to the cytoplasm (Azmi et al., 2021; Zhang et al., 2017). Here, we observed S100A10 cytoplasmic inclusions that colocalized with Crm-1 near the nucleus of atrophic astrocytes in old and aged three shrews. In aging, an enhanced Crm-1 activity alters autophagy and promotes senescence (Gorostieta-Salas et al., 2021), while the inhibition of Crm-1 expression increases longevity (Kumar et al., 2018). Here, we show increased cS100A10+ and ncS100A10+ astrocytes in old and aged subjects, which could be related to increased Crm-1 activity, and eventually with astrocytic atrophy.

Future studies should analyze other astroglial markers, such as vimentin, nestin, or S100β, to analyze how aging affects different astroglial populations during tree shrew brain aging.

## 5. Conclusion

This study demonstrated the presence of astroglial atrophy in selected brain regions of the three shrews during the course of aging. RNA-oxidation was increased in aged animals in all brain regions, which indicates that oxidative stress may not be the leading cause of astrocytic atrophy. S100A10 protein in the nucleus is associated with larger astrocytes, while nuclear S100A10 label decreases with aging. Furthermore, cytoplasmic S100A10 was found in atrophic astrocytes, in old and aged tree shrews, which suggests that loss of nuclear S100A10 during aging could be associated with the atrophic process.

## CONFLICT OF INTEREST

The authors declare no conflict of interest.

## DATA AVAILABILITY STATEMENT

The data supporting this study’s findings are available from the corresponding author upon reasonable request.

## ACKNOWLEDGMENTS

Rodriguez-Callejas, J.D. CONACYT Scholarship no. 308515.

---

**Supplementary table 1.** Specifications of primary and secondary antibodies used in the current study.

| **Primary antibodies** |  |  |  |  |
| --- | --- | --- | --- | --- |
| **Antibody** | **Host/Isotype** | **Brand** | **Catalog number** | **Dilution** |
| Anti- glial fibrillar acidic protein (GFAP) | Goat/IgG | Abcam | ab53554 | 1:300 |
| Anti-8 hydroxyguanosine (8OHG) | Mouse/IgG | Abcam | ab62623 | 1:10000 |
| Anti-S100A10 | Mouse/IgG | Invitrogen | MA-515326 | 1:250 |
| Anti - Crm-1<br>*Kindly donated by Dr. Susana Castro Obregon. | Rabbit/IgG | Novus | 100-79802 | 1:500 |
| **Secondary antibodies** |  |  |  |  |
| Cy3-conjugated anti-rabbit | Donkey/ IgG | Jackson Immuno Research | 711-165-152 | 1:500 |
| ALEXA488-conjugated anti-mouse | Donkey/IgG | Jackson Immuno Research | 715-545-150 | 1:500 |
| ALEXA647-conjugated anti-goat | Donkey/IgG | Jackson Immuno Research | 705-605-147 | 1:500 |

**Supplementary figure 1.**
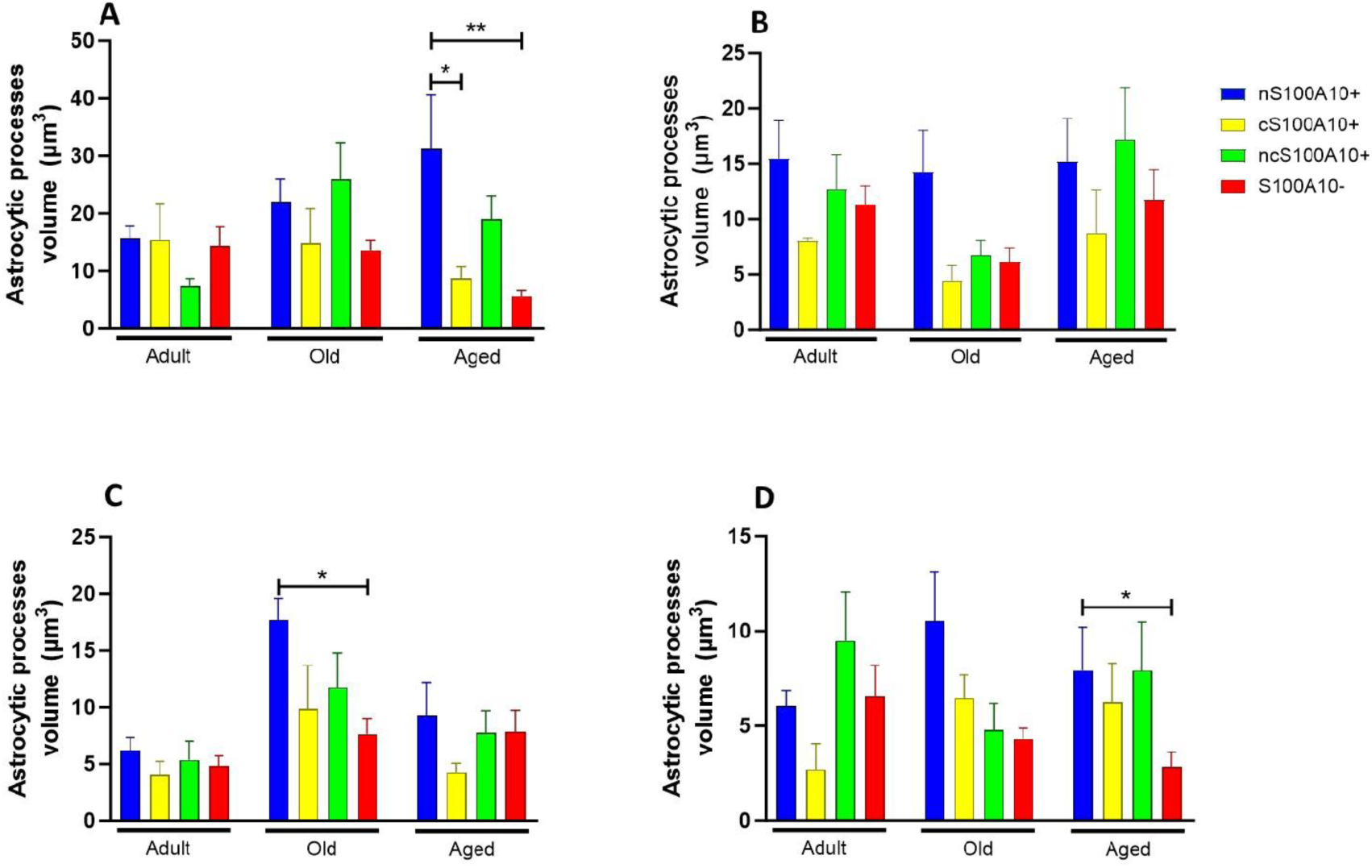
Quantification of APV in nS100A10+, cS100A10+, ncS100A10+ and S100A10- astrocytes. **(A)** DG. (**B)** CA3. (**C)** CA2-CA1. **(D)** SUB. In **adult tree shrews**, the four types of astrocytes present similar levels of APV in DG and CA2-CA1. In CA3 and SUB, the APV of cS100A10+ tends to be lower than the other types (not significant). In **old tree shrews**, nS100A10+ presents the highest levels of APV in CA3, CA2-CA1, and SUB (significant differences in CA2-CA1 nS100A10+ vs. S100A10-). In DG, the APV of the four types were similar. In **aged tree shrews**, APV of nS100A10+ were higher than the other types in DG and SUB (significant differences in DG: nS100A10+ vs. cS100A10+ and nS100A10+ vs. S100A10-; SUB: nS100A10+ vs. S100A10-). In the case of CA3 and CA2-CA1, the levels of APV in the four types were similar. Data represents means ± SEM One-way ANOVA, Tukey post hoc analysis. *p < 0.05, **p < 0.01.

## Notes

### Competing Interest Statement

The authors have declared no competing interest.

